# Acorn-omics: Optimal Foraging Behaviour Generates Steep Discounting and Preference Reversals in Laboratory Tasks

**DOI:** 10.64898/2026.07.13.738182

**Authors:** Y.L. Burnham, T.W. Fawcett, L.A. Leaver, A.D. Higginson

## Abstract

Classic foraging models and discounting tasks may oversimplify the decision-maker’s environment, resulting in a discrepancy between predicted and observed behaviour. In delay discounting tasks, animals typically steeply devalue larger-later (LL) outcomes, choosing smaller-sooner (SS) rewards after short delays. This steep discounting appears to be irreconcilable with natural foraging behaviour, where animals frequently endure long delays when travelling to find food, handling tough items, and storing food for future use. This apparent mismatch in behaviour has led to questions regarding the ecological validity of laboratory discounting tasks. Here, we developed a rich dynamic optimisation model to identify the conditions under which animals should choose LL or SS outcomes. In our model, a food-storing animal encounters food items differing in energy content and handling time and must decide which items to eat immediately and which to cache, so that it has enough stored food to survive winter. We simulated a range of environments, including laboratory conditions where foragers face negligible predation risk when searching for food and have a high probability of finding food, compared to more natural conditions where searching for food is risky, and food is harder to find. In line with preference reversals seen in the discounting literature, our model predicts that LL items should be rejected more often when the handling time for such items is increased, whereas SS items should be rejected more often when the handling time for both items is increased. Importantly, our model only predicts rejection of food items under parameter values that reflect laboratory conditions, supporting the notion that the steep devaluation of rewards seen in animals may be driven by the artificiality of traditional discounting tasks.

## Introduction

When foraging, animals make frequent intertemporal choices where outcomes differ in both their value and when they can be realised (Stevens & Stephens, 2010). It is currently unclear why animals are willing to expend considerable time and effort searching for and securing food items in their natural environments, yet show steep devaluation of rewards with their associated costs, known as discounting, in laboratory settings (Hayden, 2016).

In delay discounting tasks, animals choose between receiving a small reward after a short delay (smaller-sooner, SS) or a larger reward after a longer delay (larger-later, LL) (Mazur, 1984). As the delay to receiving the LL reward is increased, animals typically show an increased preference for the SS reward (Stevens & Stephens, 2010). However, when the delay to both rewards is increased by a fixed amount, animals typically switch their preference to the LL reward (known as a preference reversal; Ainslie, 1974), contrary to the stable preferences predicted under exponential discounting (Samuelson, 1937). For example, pigeons (*Columba livia*) favour the SS over the LL reward when the associated delays are 2 and 6 seconds, respectively, but favour the LL reward when both delays are increased by 26 seconds (Green et al., 1981).

In these delay discounting tasks, animals steeply devalue the LL reward, with rewards losing half their value on average after just a few seconds (Hayden, 2016). Yet, when foraging under natural conditions, animals regularly tolerate long delays. Perhaps the most pertinent example is food-storing behaviour, or caching, which is essential for survival and reproduction in many species (Vander Wall, 1990; Wauters et al., 1995). Food-hoarding species make frequent intertemporal decisions whether to eat or cache encountered food items, storing these items for up to months at a time for future periods of resource scarcity (Stevens & Stephens, 2008; Stevens & Stephens, 2010; Vander Wall, 1990). Opting for such long delays appears irreconcilable with the steep discounting observed in laboratory tasks, questioning to what degree these paradigms are ecologically valid (Hayden, 2016).

Similarly, there are some important discrepancies between observed foraging behaviour and the patterns of behaviour predicted by Optimal Foraging Theory. For example, the Optimal Diet Model predicts that a forager should specialise on a more profitable food type when it becomes sufficiently abundant in the environment (Charnov, 1976). However, when tested empirically, animals, including food-hoarding species (Lichti et al., 2017), tend to show partial preferences, where a food type is sometimes accepted and sometimes rejected (McNamara & Houston, 1987). Thus, in simplifying an animal’s environment, mathematical models and laboratory experiments may sometimes omit key features that are important for understanding foraging behaviour. If these models and experiments are simplified to the extent that they no longer reflect the natural decision-making environment to which animals are adapted, then the predictions may fail to align with the observed behaviour, leading us to erroneously interpret the decisions animals make as irrational (Fawcett et al., 2014).

Sometimes these discrepancies can be resolved by enriching the models with ecologically relevant details. For example, partial preferences are predicted when models are extended to account for factors such as an animal’s energy reserves and the length of the foraging period (Houston & McNamara, 1985; McNamara & Houston, 1987). In the context of intertemporal choice, there is evidence to suggest that animals are willing to tolerate longer delays in more ecologically relevant tasks, where animals make sequential or distance-based decisions more in line with their natural feeding ecology, compared to standard delay discounting paradigms (Blanchard & Hayden, 2015; Stephens & Anderson, 2001; Stevens et al., 2005a; 2005b). We might therefore question whether the high levels of rejection seen in laboratory discounting tasks reflect natural foraging behaviour or are instead a result of oversimplified environmental conditions.

To address this question, we developed a state-dependent dynamic programming model (Mangel & Clark, 1989) to determine the optimal foraging strategy for a food-hoarding individual in an environment that incorporates key ecological pressures, including predation risk, cache pilferage and variable food availability. We compare the predicted behaviour from our dynamic model to that from a static model based on net rate of gain, like the Optimal Diet Model, and to observed behaviour in delay discounting tasks, including findings on when preferences reverse. We find that including these key ecological pressures in the state-dependent model allows us to predict both the patterns of high rejection for a larger food reward with an increasing delay, and preference reversals commonly seen in laboratory discounting tasks.

## Methods

To understand how animals should value immediate and delayed outcomes, we built a state-dependent model in which foraging decisions can depend on the current socioecological conditions, including food availability, predation, and pilferage risk, the energetic value of the items a forager has in its caches, and its current internal energy reserves. At each decision point, an individual with energy reserves *x* and energetic value of items cached *c* encounters a food item of type *j*: either no food (*j* = 0), a smaller-sooner item (*j* = SS), or a larger-later item (*j* = LL) that takes longer to handle but provides more energy (**Table 8**). Although our model is devised with a scatter-hoarder in mind, which typically stores one food item per cache (Morris, 1962), our model is also applicable to larder-hoarders, which store many food items in a concentrated location (Smith & Reichman, 1984).

We modelled behaviour across autumn, where the animal must always keep *x* > 0 to avoid starvation, and have enough energy cached when autumn ends *c* ≥ *λ* to have a higher chance of surviving winter (**Table 1**). The onset of winter is stochastic: specifically, we assume that the probability *ψ* of winter starting in the current time step follows a Poisson distribution, such that the expected length of autumn is 1 / *ψ* decision points (**Table 1**). We use backwards recursion, starting from the expected future fitness of a forager at the onset of winter, and working backwards to determine the optimal decisions for each state across autumn.

**Table 1.**
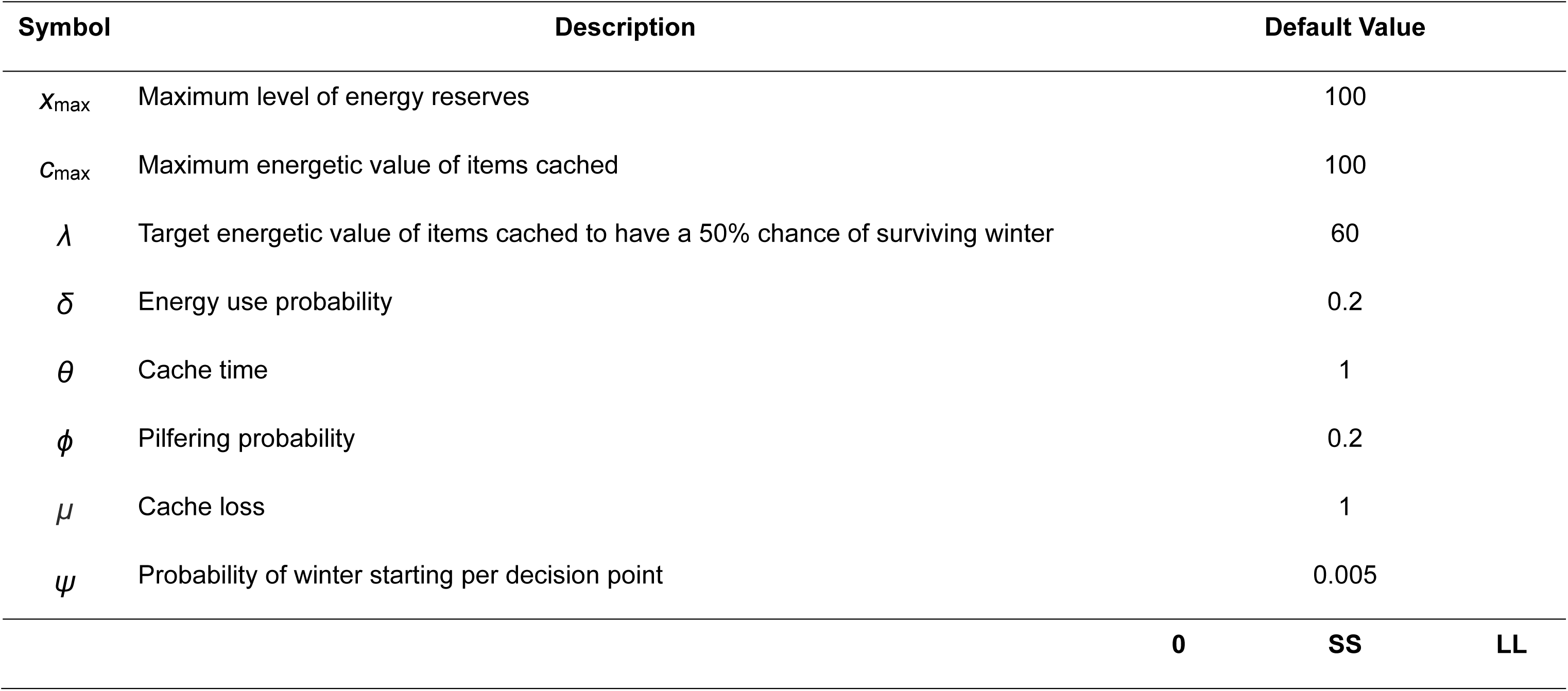
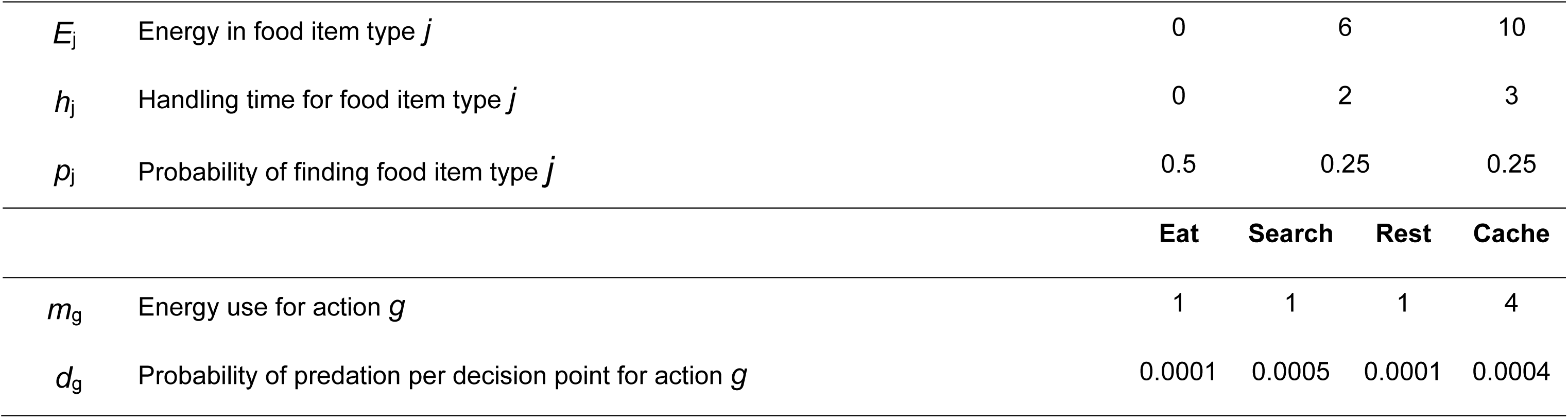
Parameters in the model and their default values.

### Parameters

### Decision-Making Process

When encountering a food item of type *j*, the forager can choose one of four actions, which we label *g*: search for a different item (*g* = search), eat the item (*g* = eat), cache the item (*g* = cache), or rest in a safe place (*g* = rest). We assume that both energy expenditure and cache loss are stochastic; when choosing action *g*, the forager uses *m*_g_ energy per unit time with probability *δ* and *m*_g_ − 1 energy with probability 1 – *δ*, and loses *μ* energy from its caches per unit time with probability *ϕ* and *μ* − 1 energy with probability 1 − *ϕ.* Since the model uses an integer state grid, and energy use was included as a non-integer, it was therefore computationally convenient to treat energy expenditure as a yes / no event occurring with a probability of 0.2. As a result, a forager’s optimal strategy depends on this probability, as it determines the expected energy use across decision points.

If the forager searches (*g* = search), the probability that it encounters a food item of type *j* before the start of the next decision point is given by *p*_j_; otherwise (*g* ≠ search), it will not find any food (*p*_0_ = 1). We calculate the possible outcomes for each action, in terms of the forager’s energy reserves, the energetic value of items stored in their caches, and the type of food item (if any) encountered before the start of the next decision point (**Fig. 1**). We denote the next states by adding a prime (*e.g.*, *x′*).

**Figure 1.**
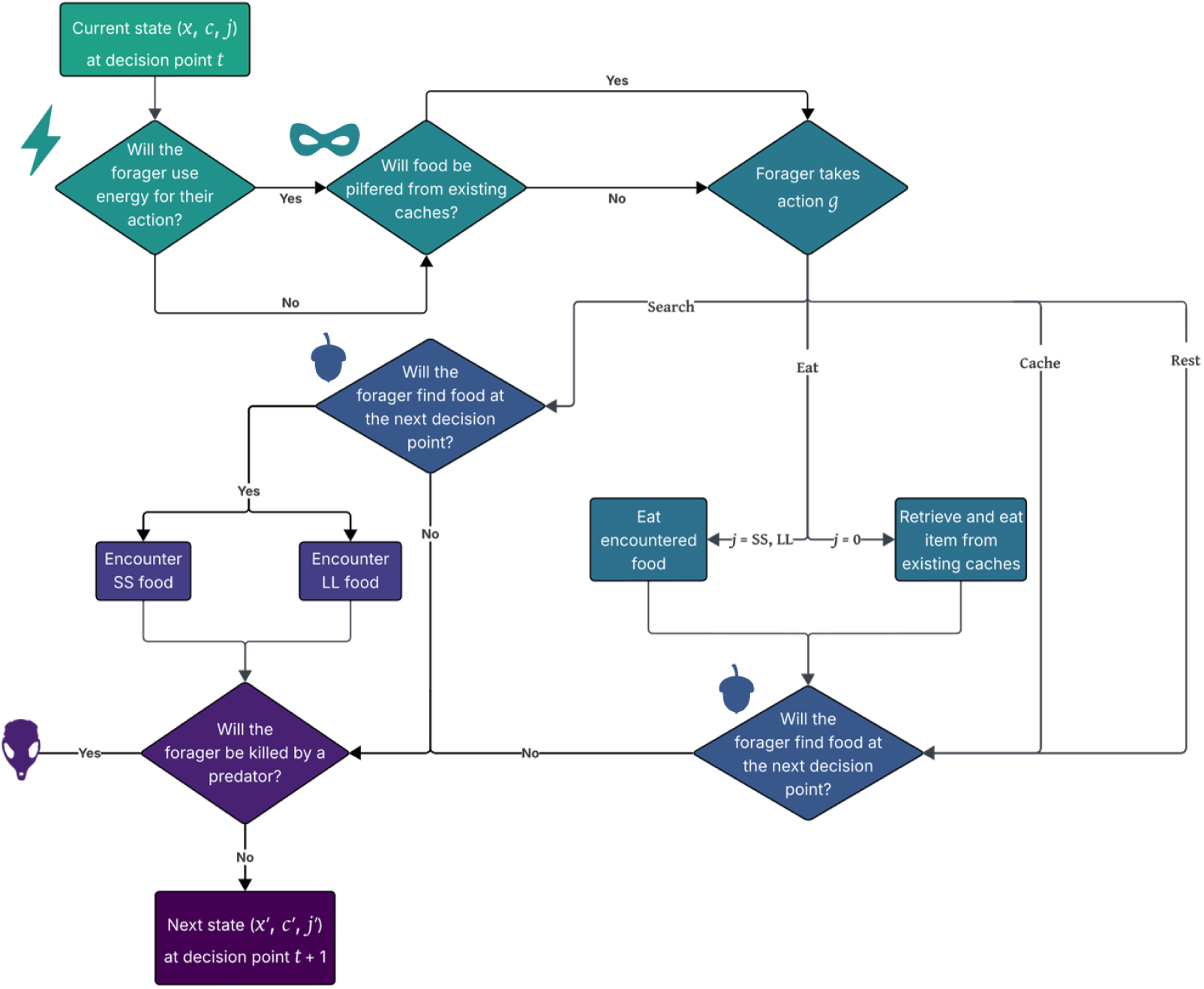
The decision-making process. We start with a forager’s current state (*x*, *c*, *j*) at the start of the current decision point, then determine how much energy the animal will use for the action it will take, and whether food will be pilfered from its existing caches. Next, the forager chooses action *g*. If it searches, it may find an SS or an LL food item at the next decision point. Finally, we determine whether the forager dies through predation. If it does not die, its next state is (*x′*, *c′*, *j′*) at the next decision point.

#### Search

When the forager fails to find a food item or rejects one, it can search for a new item. The forager’s energy reserves at the end of the decision point are calculated as:

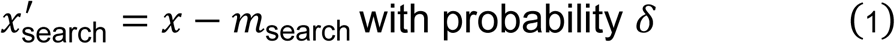

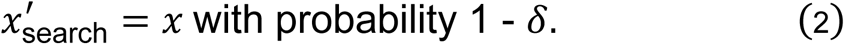

The energetic value of the items in the forager’s caches at the end of the decision point is calculated as:

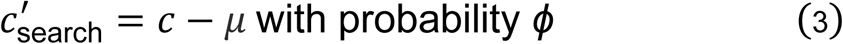

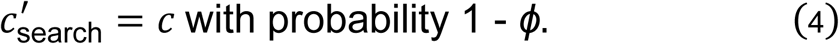

When searching, the forager encounters a food item at the next decision point with a probability of *p*_j_. The forager faces a predation risk at the current decision point of *d*_search_. We assume that the forager faces the greatest risk of predation when it is searching because its nose is to the ground, reducing the animal’s chances of detecting any potential predators (Verdolin, 2006).

#### Eat

When eating, the forager can choose to eat an encountered item or retrieve and eat from its existing caches. If the forager eats an encountered item, its energy reserves at the next decision point are calculated as:

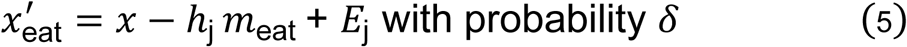

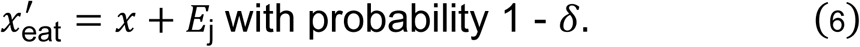

So, the forager will expend more energy eating a food item with a larger handling time.

The energetic value of the items in the forager’s caches at the next decision point becomes:

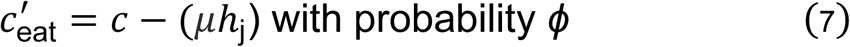

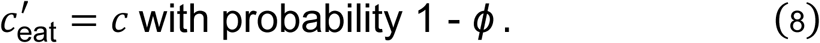

where the energy the animal loses from its existing caches depends on the time it spends handling the food item.

The forager faces a predation risk over *h*_j_ of:

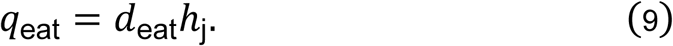

If the forager eats an item from its existing caches, that item may be an SS or an LL item, so we average these values. Its energy reserves at the end of the decision point become:

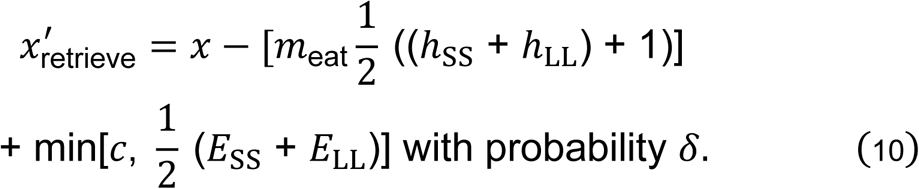

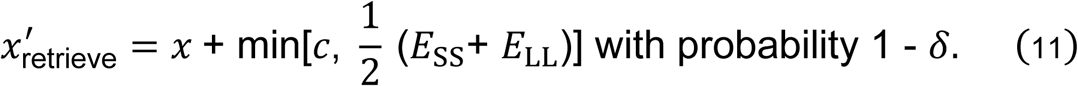

The forager gains the average energy value of the SS and LL food items (or whatever is left in its caches, if this is smaller). It pays an energy cost for the average handling time of the two food items, plus a constant time for retrieval (1), reflecting the additional time to dig up the cache. Thus, it is never beneficial for a forager to eat from their existing food stores when they have encountered an item, due to the additional time cost.

Following this, the energetic value of the forager’s caches at the next decision point becomes:

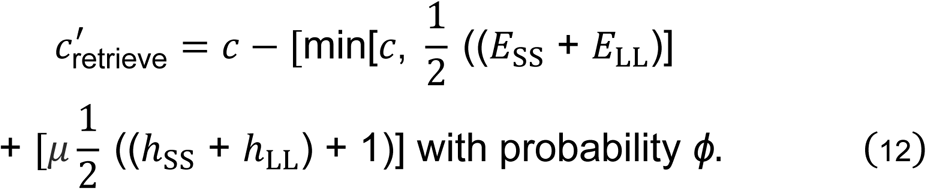

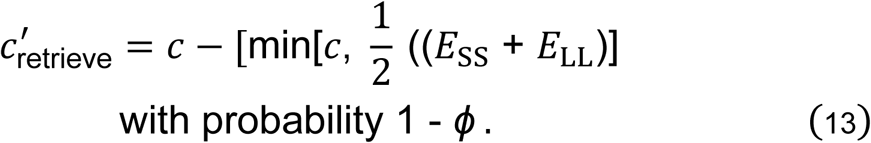

where the energy the forager loses from its existing caches scales with handling time, plus the constant time for retrieval.

Finally, the chance that the forager is killed by a predator before the next decision point is:

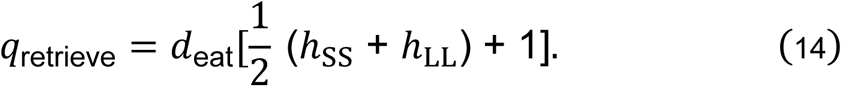

where predation risk scales with the average handling time of the two foods, plus the constant retrieval time. We assume that retrieving an item to eat it takes less time and energy than storing it, as the forager knows where to find the item, reducing time spent away from cover, and does not have to expend energy pushing the item into the ground and covering the cache.

#### Cache

When a forager decides to cache the food item it encounters, its energy reserves at the next decision point are calculated as:

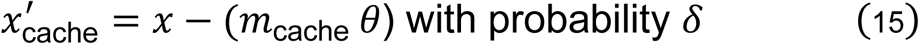

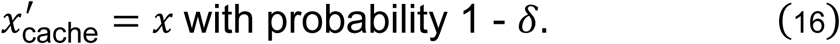

Caching is energetically costly (Helgestad et al., 2026), especially digging a hole for the item to be placed in (Vander Wall, 1993). We assume that caching is quicker than eating (Hadj-Chikh et al., 1996), and thus model this as a fixed cost, yet acknowledge that this may in reality depend on factors such as the value of the item that is being stored (Stapanian & Smith, 1978; Hopewell & Leaver, 2008), and competition (Hopewell et al., 2008).

The energetic value of the items the forager has in its caches at the next decision point becomes:

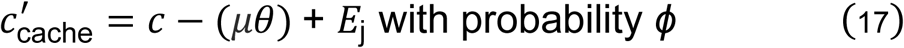

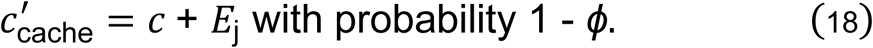

where the energy pilfered depends on the amount of time the forager spends caching the food item. Once the forager has finished caching, its existing stores are increased by the energy content of the food item.

The chance that the forager is killed by a predator before the next decision point is:

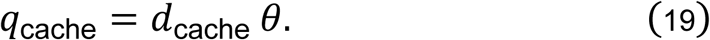

where the risk scales with how long it takes the forager to store the item. We assume that caching entails a higher predation risk than eating (*d*_cache_ = 0.0004), as animals generally move further from the safety of cover to store items, compared to foraging (Leaver et al., 2017).

#### Rest

A forager can rest to reduce its risk of being killed by a predator. Its energy reserves and the energetic value of its caches at the next decision point are calculated the same as for searching (equations (7 – 10)). The predation risk is set to the base value (*q*_rest_ = 0.0001), reflecting that resting is the safest option.

Temporal discounting may lead a forager to prioritise more immediate outcomes, such as retrieving and eating cached items before the onset of winter and eating rather than caching an encountered item. Although the process of caching an item is quicker than eating one in our model, if the animal eats an SS item immediately it pays a time cost of *h*_SS_, whereas if it caches and later retrieves the item, it pays a time cost of *θ* + (*h*_SS_ + *h*_LL_) / 2 + 1, meaning that caching still results in a more delayed outcome compared to eating.

### Fitness Functions

Fitness outcomes in the model are based on the chance that the forager survives winter (**Fig. 2**). At the onset of winter, fitness is specified by a terminal fitness function (*u_c_*) based on the energetic value of the items the forager currently has in its caches (*c*).

**Figure 2.**
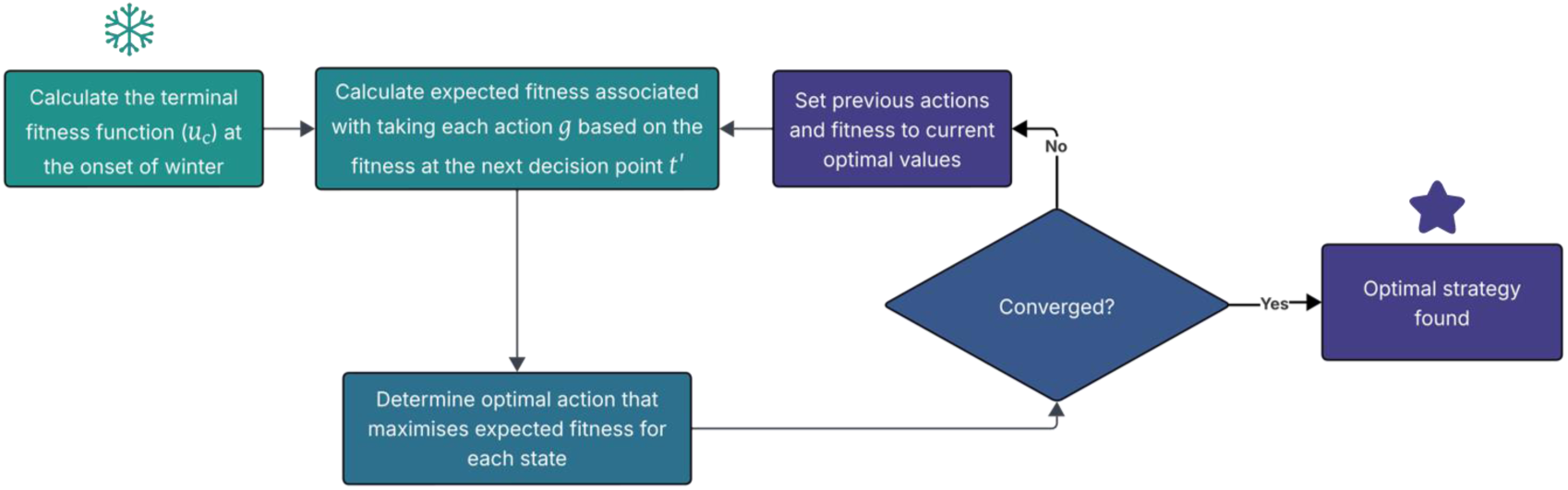
The backwards recursion process. At the onset of winter, the terminal fitness function (*u*_c_) is calculated, defining the fitness for each state (*x*, *c*, *j*). From here, we iterate backwards, where for any given current state at decision point *t*, we determine the expected fitness associated with taking each action *g* as a function of the fitness at the next decision point *t′*, providing us the optimal action which maximises expected fitness at that decision point. This is repeated until convergence of the optimal decisions and fitness arrays, where we have then found the optimal strategy.

*u*_c_ is calculated for each possible state, and is always a value between 0 and 1:

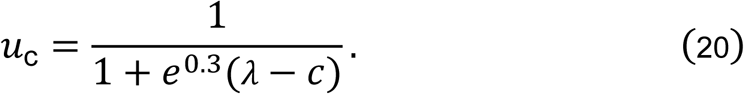

If *x* ≤ 0, *u*_c_ is set to 0, indicating that the forager has starved to death. The coefficient (0.3) determines how quickly *u*_c_ moves towards 0 or 1, dependent on whether *c* < *λ* or *c* > *λ*. For example, a forager has a 99% chance of surviving winter when the items it has cached have an energetic value of 75.

Once *u*_c_ has been calculated, the model iterates backwards, evaluating the fitness consequences of each action *g* for a forager in each state (*x*, *c*, *j*). If the forager chooses the optimal action (*i.e.* the one that maximises its expected fitness), its expected fitness is given by:

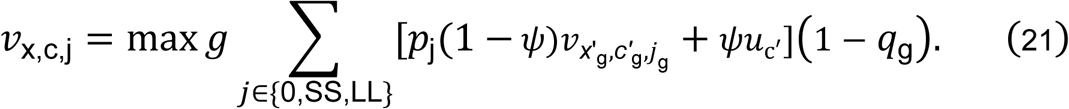

For each action *g*, the expected fitness is calculated by summing over all the possible outcomes. For each outcome, we consider both the fitness *v*_x,c,j_ if autumn continues (which happens with probability 1 − *ψ*) and the fitness *u*_c_ if winter starts (which happens with probability *ψ*). We then identify the action *g* that provides the highest expected fitness for each state (**Fig. 2**).

Finally, the model checks for convergence (**Fig. 2**). In our model, the convergence criteria are that the sum of the differences in fitness and action values (*e.g.*, search = 1, eat = 2, *etc.*) between the current and previous iteration are < 0.01 and < 1, respectively. Once the convergence criteria are met, we have found the optimal strategy (**Fig. 2**). If the convergence criteria are not met, *v*_x,c,j_ is updated with the optimal fitness values from the current iteration and a new iteration begins (**Fig. 2**). For the first 10 iterations, *v*_x,c,j_ is always updated with these optimal values to give some time for stabilisation.

### Simulation

To predict foraging behaviour, we simulated the actions and fates of 200 individuals following the optimal strategy. We started each forager with energy reserves (*x*) randomly sampled from a normal distribution and bounded between 10 and *x*_max_, no stored items (*c* = 0), and no encountered food (*p*_0_ = 1). We identified the forager’s optimal decision given its current state, then stochastically determined its next state based on the associated probabilities, iterating forward until the onset of winter occurs for most foragers, where we then calculated the forager’s chance of surviving the winter.

### Urban *vs.* Rural *vs.* Laboratory Environments

We simulated four different environmental conditions in our model. The optimal strategy was adapted to these conditions, where we assume that foragers learn from the new environment. In our urban condition, pilfer rate and predation risk are high (*ϕ* = 0.4, *d*_search_ = 0.0005), and the probability of finding food is low (*p*_SS_ = 0.25, *p*_LL_ = 0.25) (**Table 9**). In contrast, our rural condition has the same predation risk (*d*_search_) and food availability (*p*_j_), but a lower pilfer rate (*ϕ* = 0.1). We assumed that pilferage risk is higher in urban environments, due to a higher abundance of food-storing individuals (Dittel & Vander Wall, 2018). Finally, we simulated two laboratory conditions. In the first condition (laboratory with pilfering), we assumed that the animal’s cage is regularly cleared of uneaten food items, analogous to a high pilfer rate (*ϕ* = 0.4), and the animal is guaranteed to find food (*p*_SS_ = *p*_LL_ = 0.5), while predation risk is low (*d*_search_ = 0.0002). In the second condition (laboratory without pilfering), the animals’ cage is not cleaned of any uneaten food, resulting in the same *p*_j_ and *d*_g_ but a lower pilfer rate (*ϕ* = 0.1). These parameter values reflect how the forager is assumed to perceive its environment, hence there is a small predation risk and pilfer rate for the laboratory condition without pilfering, despite no actual risk of being attacked by a predator or losing stored items.

### Net Rate of Gain

To make comparisons to the Optimal Diet Model, we varied *p*_LL_ within our four environments (**Table 2**).

**Table 2.**
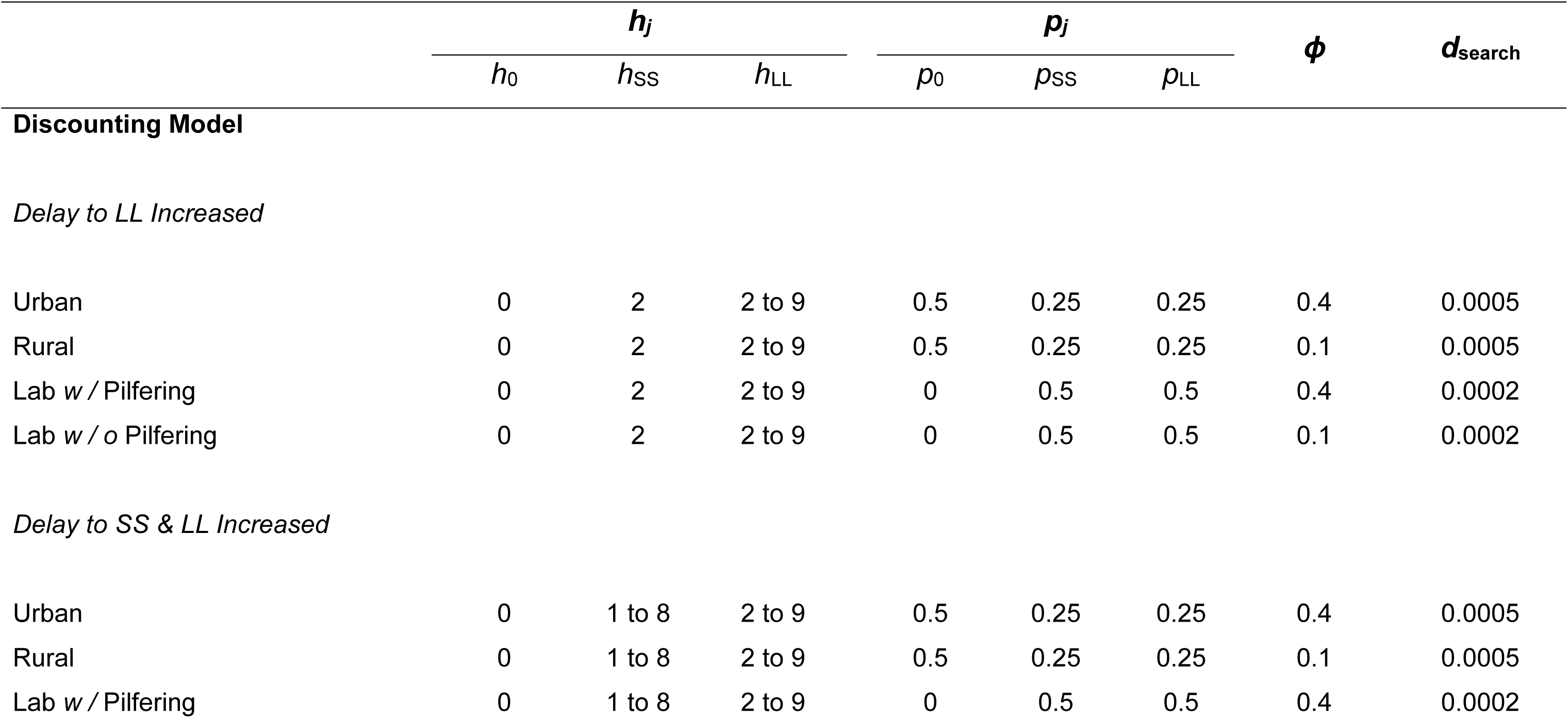
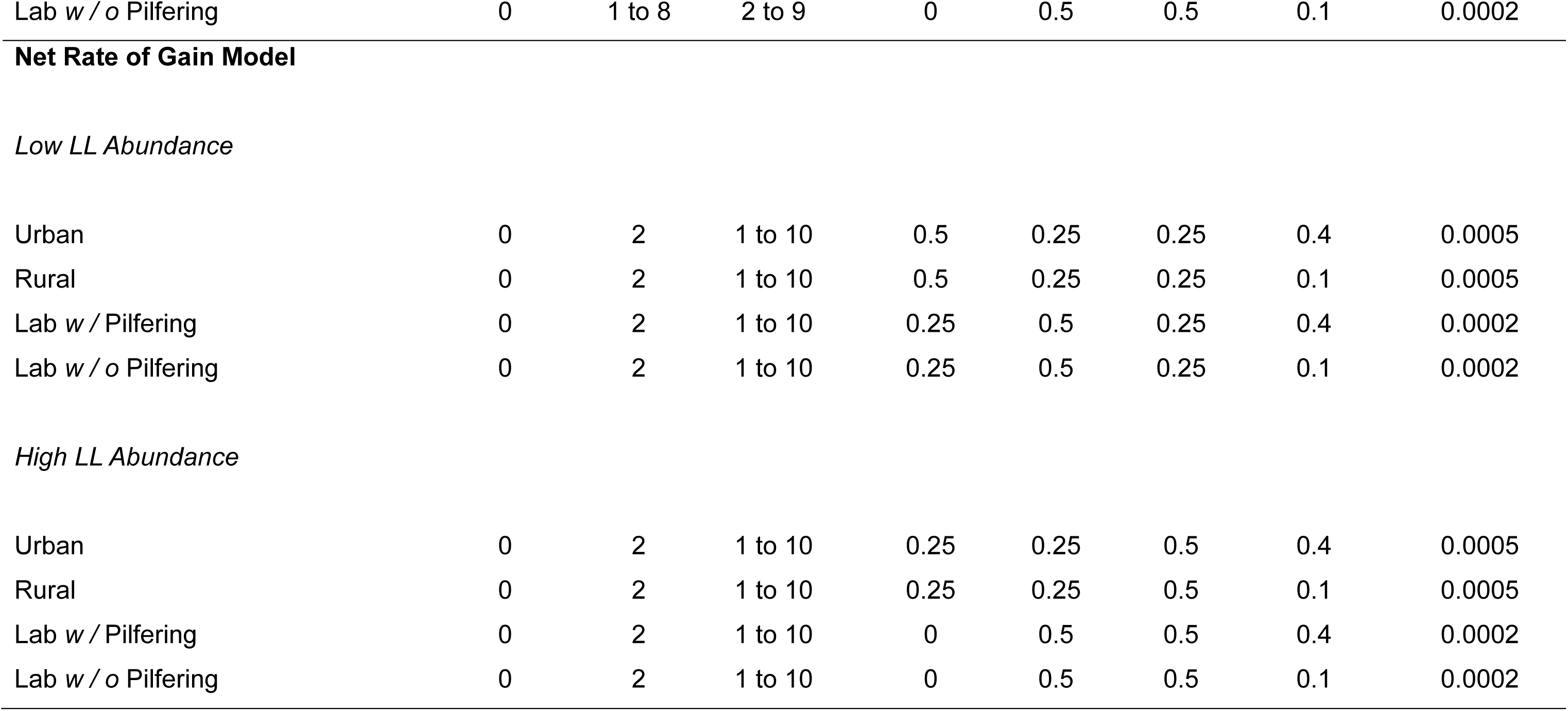
Parameter values in the models under a.) Urban conditions, b.) Rural conditions, c.) Laboratory conditions with pilfering, and d.) Laboratory conditions without pilfering when the delay to just the LL item is increased, and the delay to both the SS and LL items is increased (discounting), and during low and high LL abundance (model based on the Optimal Diet Model).

| | $h_j$ | | | $p_j$ | | | $\phi$ | $d_{\text{search}}$ |
| --- | --- | --- | --- | --- | --- | --- | --- | --- |
| | $h_0$ | $h_{\text{SS}}$ | $h_{\text{LL}}$ | $p_0$ | $p_{\text{SS}}$ | $p_{\text{LL}}$ | | |
| Discounting Model |  |  |  |  |  |  |  |  |
| Delay to LL Increased |  |  |  |  |  |  |  |  |
| Urban | 0 | 2 | 2 to 9 | 0.5 | 0.25 | 0.25 | 0.4 | 0.0005 |
| Rural | 0 | 2 | 2 to 9 | 0.5 | 0.25 | 0.25 | 0.1 | 0.0005 |
| Lab w / Pilfering | 0 | 2 | 2 to 9 | 0 | 0.5 | 0.5 | 0.4 | 0.0002 |
| Lab w / o Pilfering | 0 | 2 | 2 to 9 | 0 | 0.5 | 0.5 | 0.1 | 0.0002 |
| Delay to SS & LL Increased |  |  |  |  |  |  |  |  |
| Urban | 0 | 1 to 8 | 2 to 9 | 0.5 | 0.25 | 0.25 | 0.4 | 0.0005 |
| Rural | 0 | 1 to 8 | 2 to 9 | 0.5 | 0.25 | 0.25 | 0.1 | 0.0005 |
| Lab w / Pilfering | 0 | 1 to 8 | 2 to 9 | 0 | 0.5 | 0.5 | 0.4 | 0.0002 |
| Lab w / o Pilfering | 0 | 1 to 8 | 2 to 9 | 0 | 0.5 | 0.5 | 0.1 | 0.0002 |
| Urban | 0 | 2 | 1 to 10 | 0.5 | 0.25 | 0.25 | 0.4 | 0.0005 |
| Rural | 0 | 2 | 1 to 10 | 0.5 | 0.25 | 0.25 | 0.1 | 0.0005 |
| Lab w / Pilfering | 0 | 2 | 1 to 10 | 0.25 | 0.5 | 0.25 | 0.4 | 0.0002 |
| Lab w / o Pilfering | 0 | 2 | 1 to 10 | 0.25 | 0.5 | 0.25 | 0.1 | 0.0002 |
| *High LL Abundance* |  |  |  |  |  |  |  |  |
| Urban | 0 | 2 | 1 to 10 | 0.25 | 0.25 | 0.5 | 0.4 | 0.0005 |
| Rural | 0 | 2 | 1 to 10 | 0.25 | 0.25 | 0.5 | 0.1 | 0.0005 |
| Lab w / Pilfering | 0 | 2 | 1 to 10 | 0 | 0.5 | 0.5 | 0.4 | 0.0002 |
| Lab w / o Pilfering | 0 | 2 | 1 to 10 | 0 | 0.5 | 0.5 | 0.1 | 0.0002 |

The net rate of energy gain for being a generalist (accepting both SS and LL items) is:

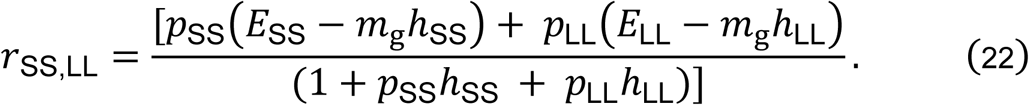

The net rate of gain for specialising on SS items is:

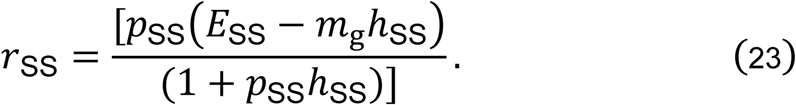

Finally, the net rate of gain for specialising on LL items is:

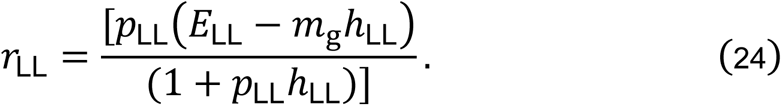

The critical handling time for the LL item, where generalising and specialising on the LL item yield the same net rate of energy gain, is:

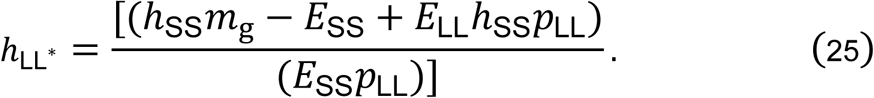

When *h*_LL_ > *h*_LL_*\**, the forager should generalise, accepting both SS and LL food items. When *h*_LL_ < *h*_LL_*\**, the forager should specialise on LL items.

The critical handling time for the LL item, where generalising and specialising on the SS item yield the same net rate of energy gain, is:

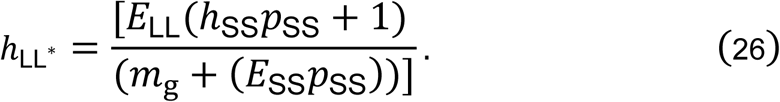

When *h*_LL_ > *h*_LL_*\**, the forager should specialise on the SS item. When *h*_LL_ < *h*_LL_*\**, the forager should generalise.

Finally, the critical handling time for the LL item, where specialising on the LL and specialising on the SS item yield the same net rate of energy gain, is:

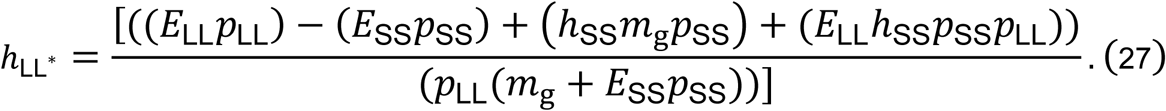

When *h*_LL_ > *h*_LL_*\**, the forager should specialise on the SS item. When *h*_LL_ < *h*_LL_*\**, the forager should specialise on the LL item.

### The Effect of Hoarding

To investigate whether the outcomes of our model explicitly rely on food-storing, we further simulated the same four environments as described above, but further restricted the forager from caching. To do this, we made the energetic gain from retrieving an item from a cache, and the increase in the energy of cached items when storing food, zero, effectively providing no benefit of hoarding behaviour.

## Results

### Optimal Strategy

Our model predicts that a forager will keep searching until it finds food, unless (a) it has a large amount of energy cached and at least a small amount of energy reserves, in which case it will rest to avoid predation; or (b) it has critically low energy reserves, in which case it will retrieve and eat an item from its stores early (**Fig. 3a**). If a forager encounters an SS or an LL food item it will cache the item unless its energy reserves are very low, in which case it will eat the item, or unless it has a large amount of energy stored, in which case it will rest to avoid predation if its energy reserves are high, and eat the item otherwise (**Fig. 3b, c**). The LL food item is cached across more of the state space than the SS food; energy reserves must be lower for the forager to eat rather than cache an LL compared to an SS item (cf. **Fig. 3b, c**; **Fig. 3d**). Under the baseline parameter settings, animals never search for another item once they have found an SS or an LL item (**Fig. 3b, c**). Foragers are most likely to have a high amount of energy stored and low energy reserves under baseline values (**Fig. 4**).

**Figure 3.**
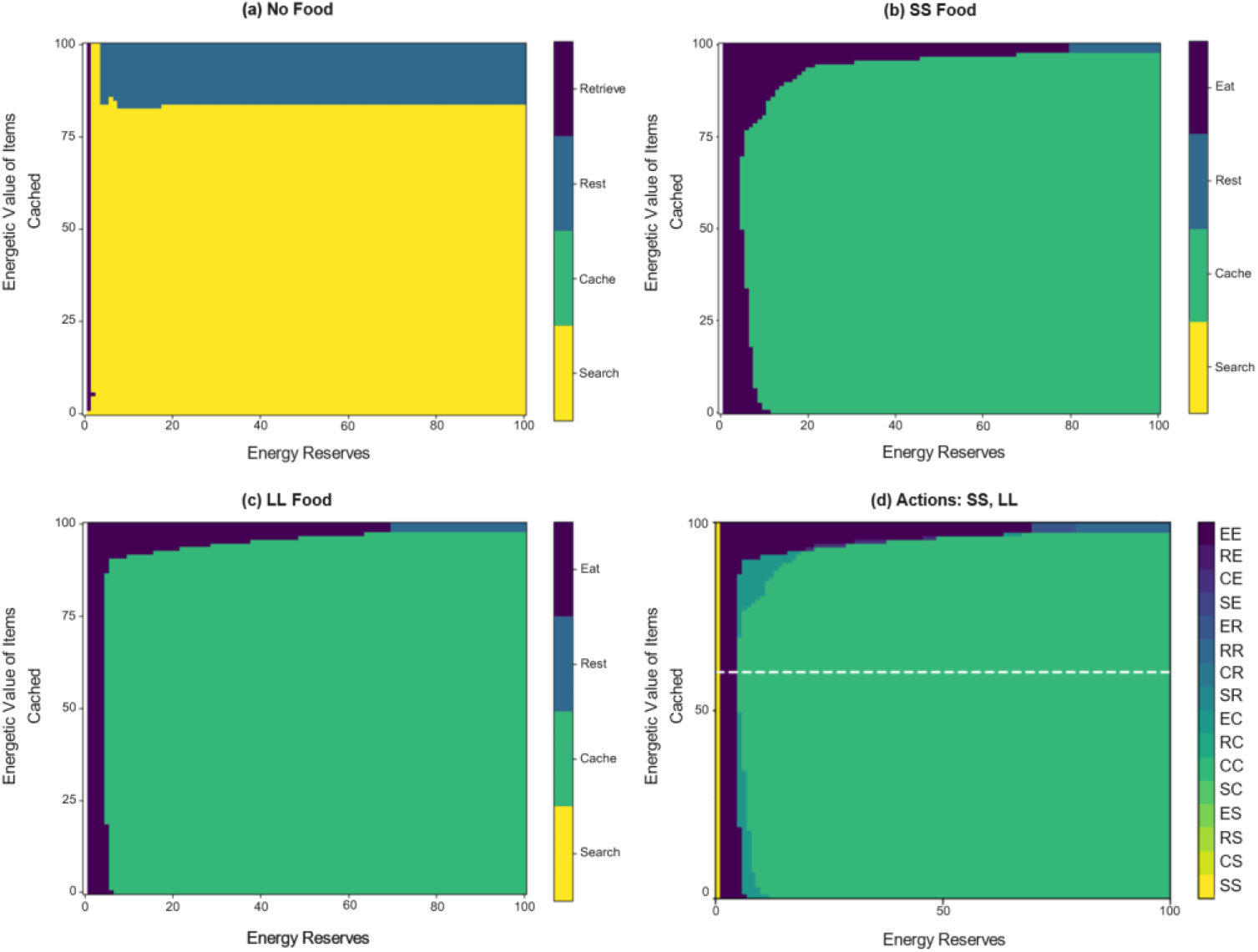
Optimal strategy dependent on the forager’s energy reserves, the energetic value of the items they have stored, and whether it encounters (a) no food, (b) an SS item, or (c) an LL item under base values (**Table 1**). (d) shows the optimal strategy for both food types, with the first letter referring to the optimal action when encountering an SS item, and the second referring to the optimal action when encountering an LL item (*e.g.*, SC = search if encounter an SS item, cache if encounter an LL item). The white dashed line illustrates *λ*.

**Figure 4.**
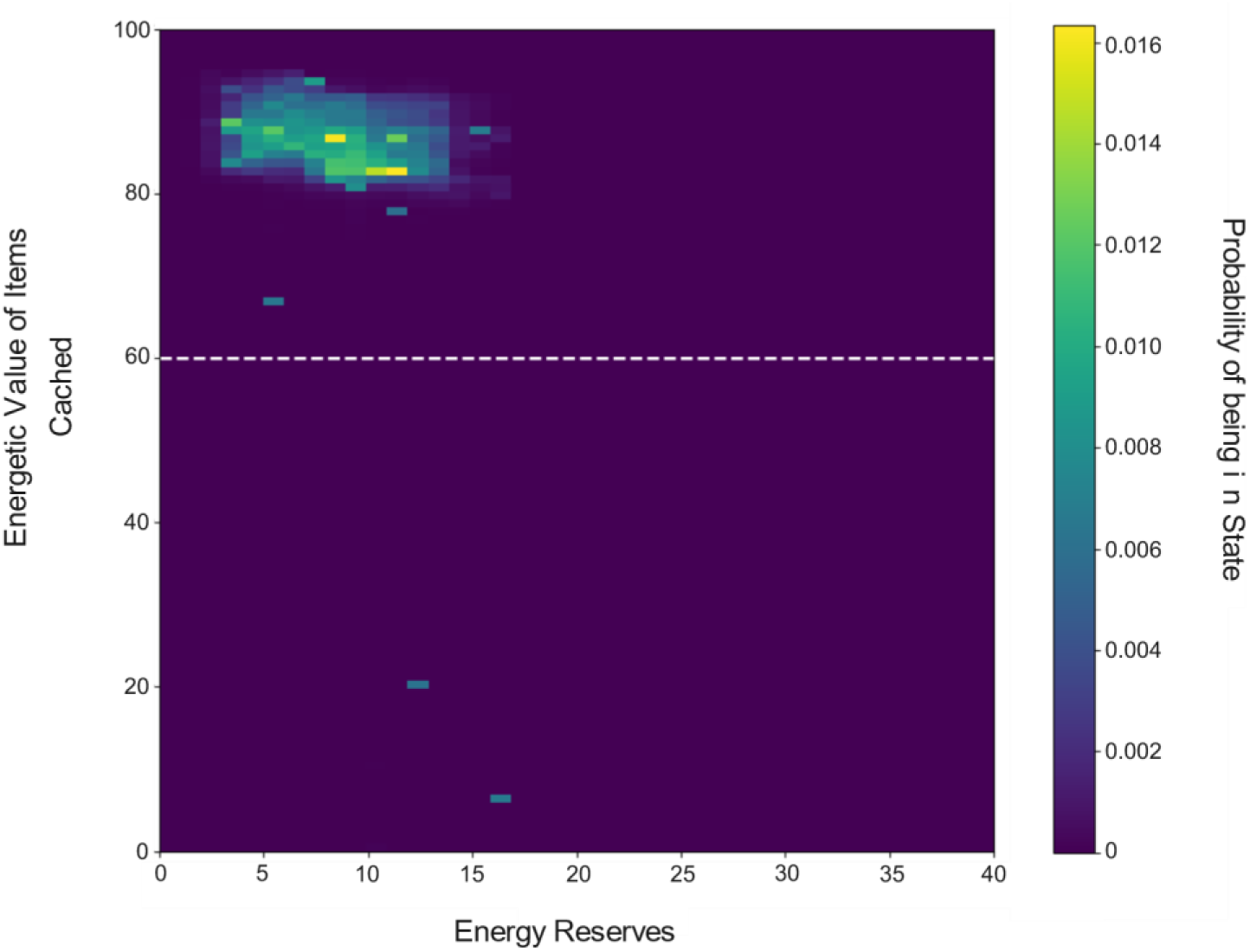
Heatmap of probabilities of forager states aggregated across all time points in the simulation under baseline values (**Table 1**). The white dashed line illustrates *λ*.

Under baseline values, when *p*_LL_ (and as a result *p*_0_) is manipulated, while *p*_SS_ is held constant, the majority of foragers survive winter (**Fig. 5**). Foragers never starve and only die through predation (**Fig. 5**). As *p*_LL_ increases, there is generally a slight increase in the number of foragers that survive winter, and a slight decrease in the number of foragers that are killed by a predator (**Fig. 5**).

**Figure 5.**
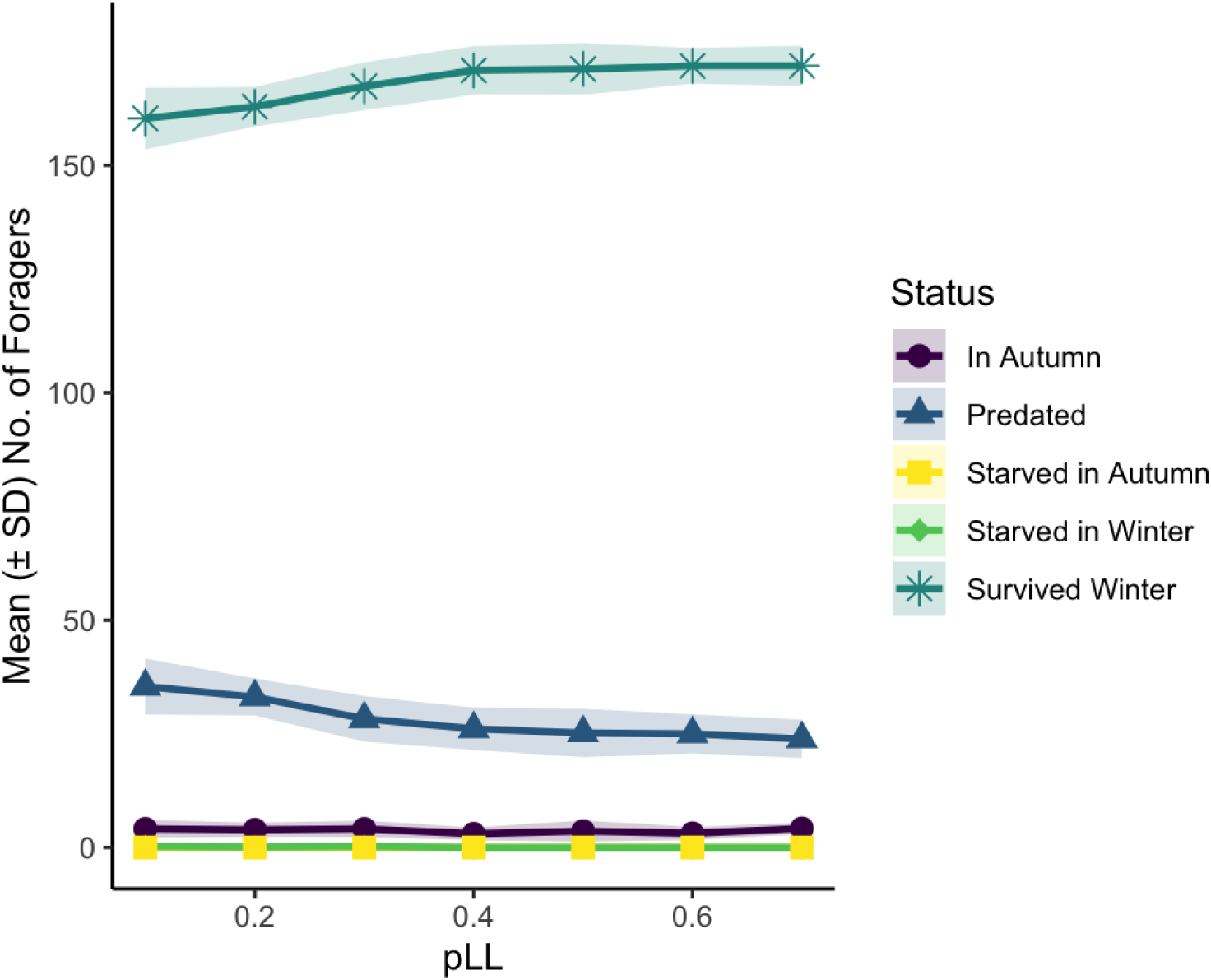
Number of foragers that starved during autumn, were killed by a predator, survived winter, starved during winter, and are alive in autumn when *p*_LL_ is manipulated, *p*_SS_ = 0.25 and *p*_0_ = 1 – (*p*_SS_ *+ p*_LL_). All other parameters assume baseline values, outlined in **Table 1**. Values represent the means (± SD) across 10 simulation runs (*N* = 200 foragers per run).

### Urban *vs.* Rural *vs.* Laboratory Environments

#### Net Rate of Gain

Predictions from our net-rate-of-gain model, based on the Optimal Diet Model, show that the forager should specialise on the SS food when the handling time for LL items is high (*h*_LL_ ≥ 7) (**Fig. 6**). In contrast, the forager should specialise on the LL food when it becomes more abundant (*p*_LL_ > 0.5) and the time to handle it is low (*h*_LL_ ≤ 2) (**Fig. 6**).

**Figure 6.**
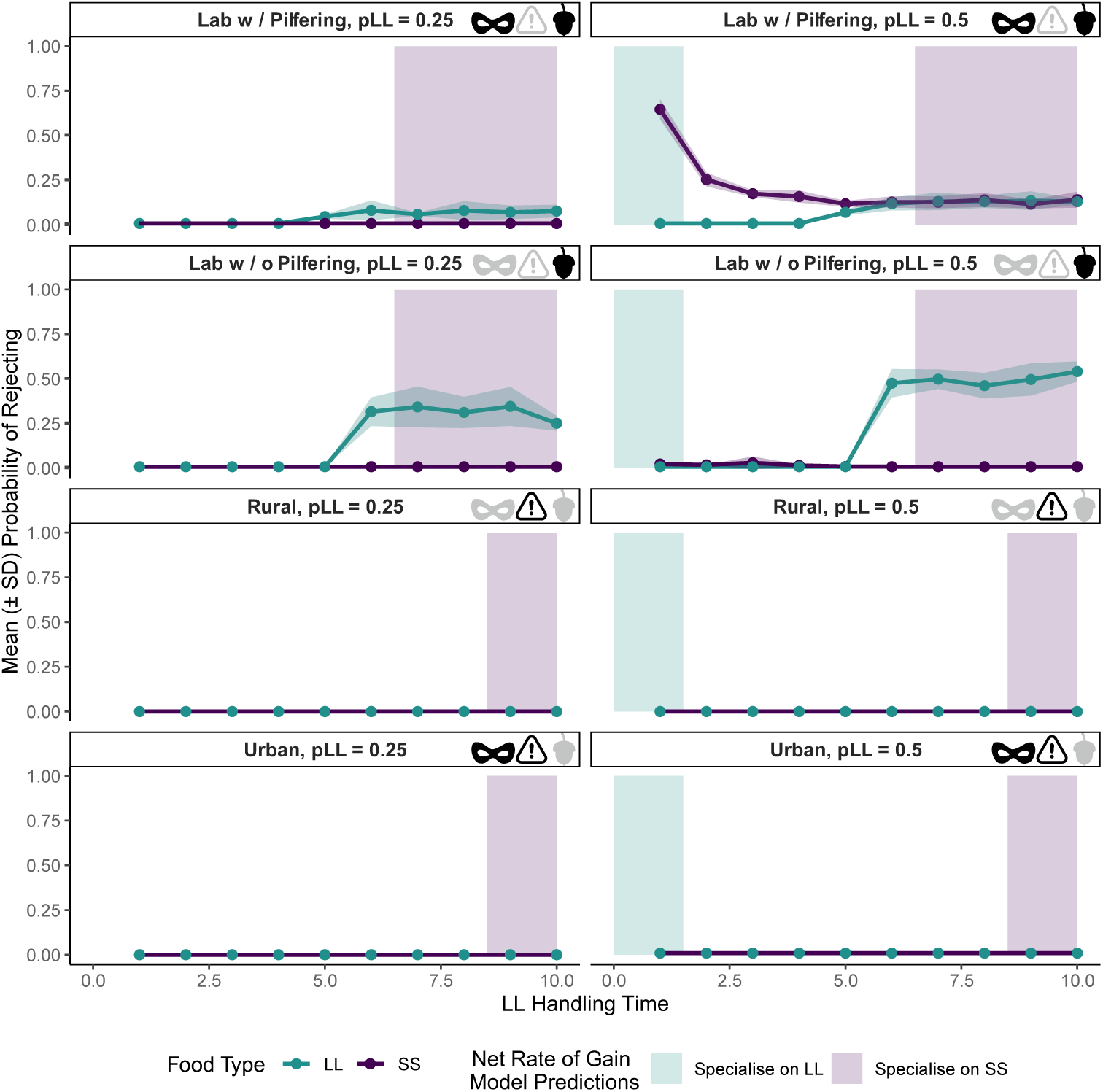
Probability of rejecting SS and LL food items, dependent on handling time for the LL item, and the probability of encountering an LL item (*p*_LL_). Predictions from the net rate of gain model are represented with the green zone illustrating when a forager should specialise on LL items, and the purple zone displaying when they should specialise on SS items. Values represent the means (± SD) across 10 simulation runs (*N =* 200 foragers per run). The parameter values are outlined in **Table 2**. The icons represent, in order, the pilferage risk (eye mask), predation risk while searching (warning sign), and probability of finding an SS item (acorn) associated with the environment, with black icons indicating these risks / probabilities are high, and grey icons indicating they are low.

In contrast to predictions from the net-rate-of-gain model, the predictions from our state-dependent model suggest that a forager should never exclusively specialise on SS or LL food items, and never reject either item in urban or rural environments (**Fig. 6**). A forager may reject an SS food item under laboratory conditions with pilfering, when the handling time for the LL item is low. A forager may reject an LL food item under laboratory conditions with or without pilfering, when the handling time for this item becomes too large (**Fig. 6**).

#### Discounting

##### Probability of Eating *vs.* Caching

When the handling time for the LL food *h*_LL_ is increased while *h*_SS_ is held constant, foragers are always more likely to eat the SS food and cache the LL food (left-hand panels; **Fig. 7**). The probability of eating the LL food tends to decrease as its handling time increases, whereas the probability of eating the SS food increases. Foragers are least likely to eat the LL food in the scenarios with high pilferage (laboratory with pilfering, and urban). In contrast, when the handling time for both the SS and the LL foods is increased simultaneously by the same amount, the probability of eating the SS food falls, whereas the probability of eating the LL food increases (right-hand panels; **Fig. 7**). At long handling times, we see a reversal where the forager’s tendency to immediately eat rather than cache encountered food is stronger for LL items than SS items. This pattern is most prominent in the laboratory scenarios, where predation risk while searching is low, and the probability of finding food is high.

**Figure 7.**
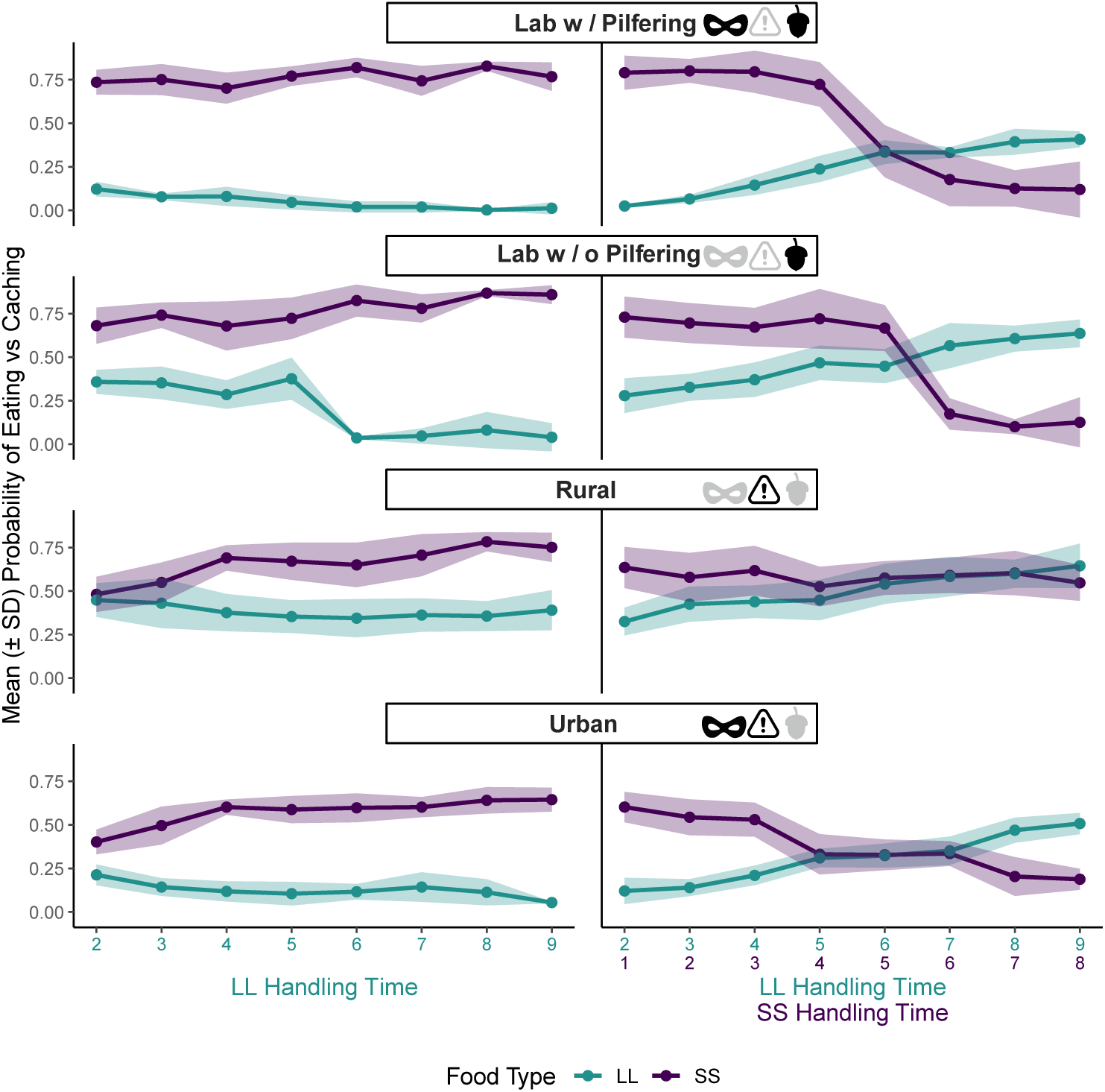
Probability of eating *vs.* caching SS and LL food items in urban, rural, and laboratory environments. Values represent the means (± SD) across 10 forward simulation runs (*N* = 200 foragers per run). The parameter values are outlined in **Table 2**. The icons represent, in order, the pilferage risk (eye mask), predation risk while searching (warning sign), and probability of finding an SS item (acorn) associated with the environment, with black icons indicating these risks / probabilities are high, and grey icons indicating they are low.

##### Probability of Rejecting Food

Our model predicts that foragers will only reject food items and search for other food items in laboratory scenarios, when predation risk while searching is low (*d*_search_ = 0.0002) and the probability of finding food is high (*p*_SS_ = *p*_LL_ = 0.5) (**Fig. 8**); in urban and rural settings, they accept all encountered items. When the handling time for the LL food *h*_LL_ is increased while *h*_SS_ is held constant, foragers begin to reject LL items in laboratory conditions. When the handling time for both foods is increased by the same amount, we see high rejection for the SS food if the time costs are so high that the forager is likely to find an LL item in this period.

**Figure 8.**
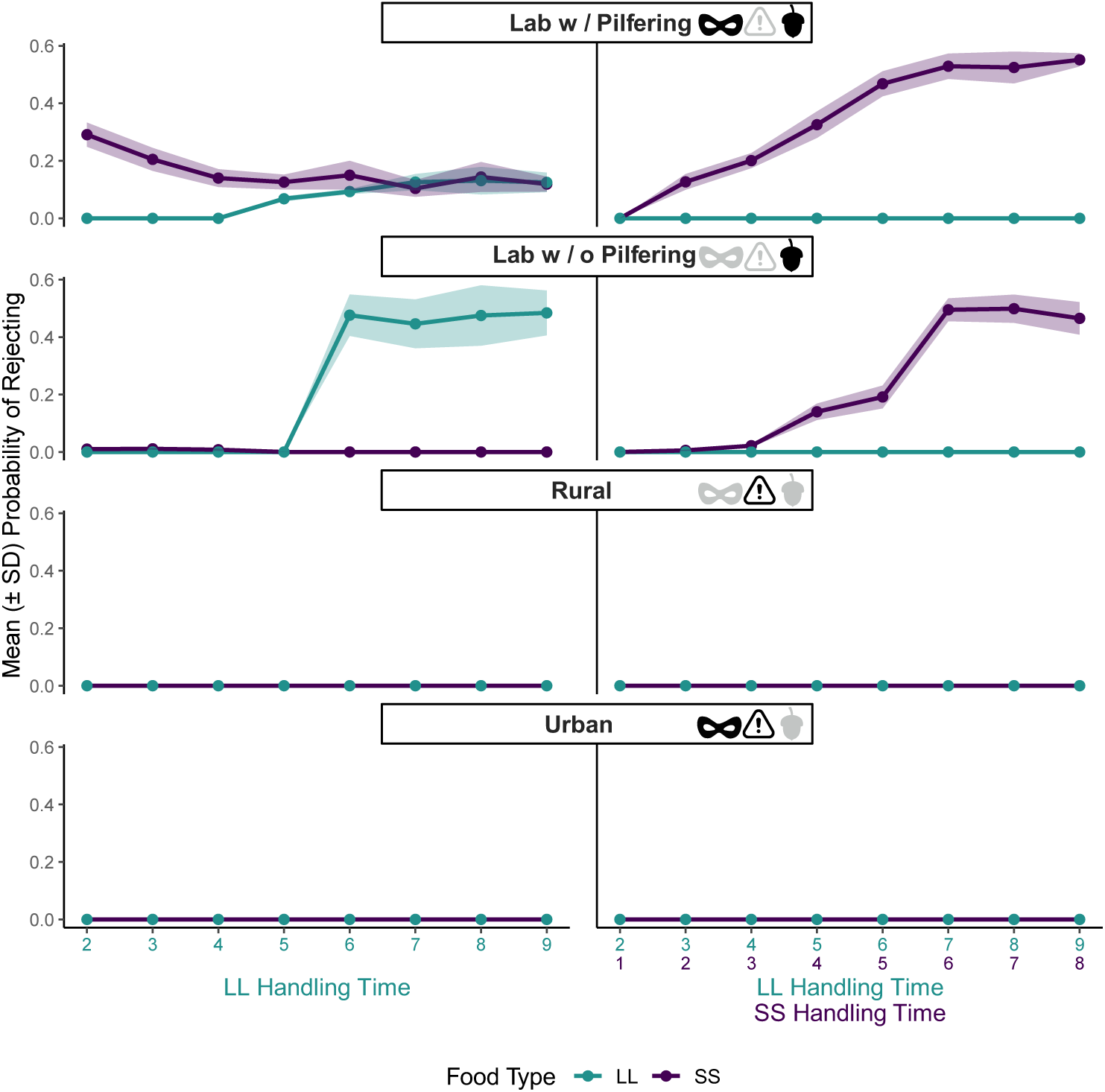
Probability of rejecting SS and LL food items in urban, rural, and laboratory environments. Values represent the means (± SD) across 10 simulation runs (*N* = 200 foragers per run). The parameter values are outlined in **Table 2**. The icons represent, in order, the pilferage risk (eye mask), predation risk while searching (warning sign), and probability of finding an SS item (acorn) associated with the environment, with black icons indicating these risks / probabilities are high, and grey icons indicating they are low.

##### The Effect of Hoarding

For non-hoarding foragers, our model predicts the same influence of environment on rejection behaviour; foragers rarely ever reject food items in rural and urban environments (**Fig. 9**), yet show steep rejection for LL foods when the handling time is increased to this food item only, and steep rejection for SS foods when the handling time to both food items is increased in laboratory environments (**Fig. 9**).

**Figure 9.**
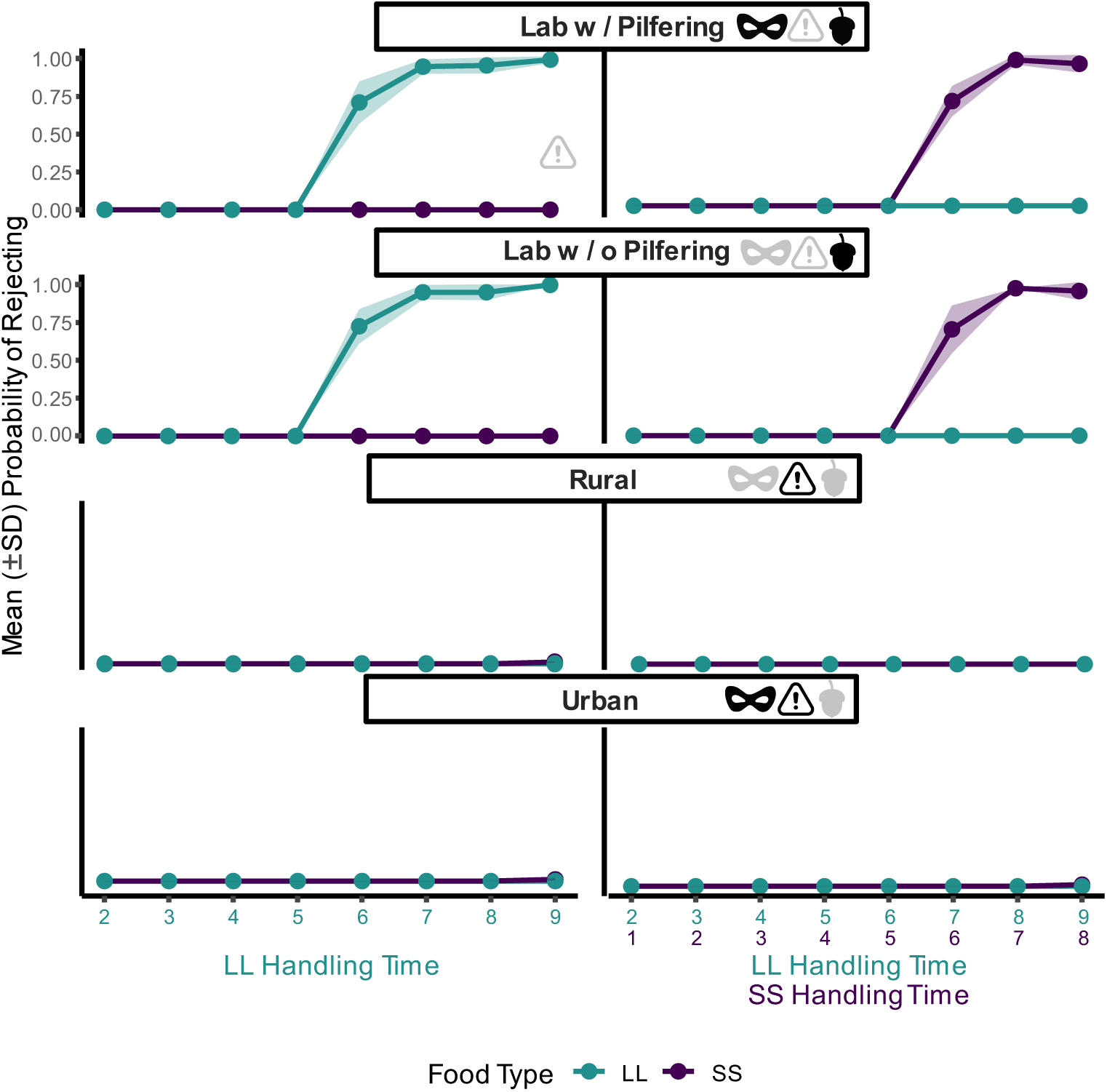
Probability of rejecting SS and LL food items in urban, rural, and laboratory environments for a non-hoarding forager. Values represent the means (± SD) across 10 simulation runs (*N* = 200 foragers per run). The parameter values are outlined in **Table 2**. The icons represent, in order, the pilferage risk (eye mask), predation risk while searching (warning sign), and probability of finding an SS item (acorn) associated with the environment, with black icons indicating these risks / probabilities are high, and grey icons indicating they are low.

## Discussion

In our model, food-storing animals are predicted to eat or retrieve items from their caches when they are at a higher risk of starvation. Foragers never rejected an encountered food item in simulated natural environments. When the risk of not finding another food item and falling to predation while searching is reduced under laboratory conditions, foragers are predicted to increase rejection of LL food as the handling time for such items is increased, and to increase rejection of SS food as the handling time for both items is increased. Partial preferences are always observed; a forager is never predicted to specialise exclusively on either food type in our model.

### Risk of Starvation

Under the optimal strategy, foragers prioritise more immediate outcomes, such as eating encountered items as soon as they find them or retrieving stored items from their caches, when they have low energy reserves. This is to reduce the risk of starvation, akin to Houston and McNamara’s (1985) model, where animals with low energy reserves are predicted to be more willing to accept less profitable prey types, prioritising immediate survival over long-term rate of gain. This prediction is supported by observations that food-storing animals will switch from eating to caching when they are satiated (Cristol, 2001; Jacobs, 1992), and by the discounting literature, where food-deprived animals are less willing than non-deprived animals to wait for a larger delayed reward, opting for more immediate outcomes (Bateson et al., 2015; Mayack & Naug, 2015; Snyderman, 1983).

In our model, foragers tolerated lower energy reserves before switching from caching to eating LL items, compared to the equivalent switch for SS items. When a forager’s energy reserves fall, they do not abandon caching altogether, but instead stop caching SS items first, and only stop caching LL items if their energy reserves drop even further. Storing LL items provides a higher future payoff because these items contain more energy than SS items. In line with this, it has been argued that food-storing animals should be more likely to eat lower-quality food items when hungry, and cache higher-quality items when satiated (Preston & Jacobs, 2009).

When we increased handling time for the LL food in our model, foragers became more likely to eat SS items. This may compensate for a decrease in the probability of consuming LL items due to a reduced net gain. The high handling time of the LL food makes it less efficient for foragers to eat, rather than cache, these items, so the animal instead opts for immediately consuming SS items to increase their energy reserves. This prediction is supported by empirical work on grey squirrels (*Sciurus carolinensis*), which preferentially cache food items with high handling times, such as chow blocks and unshelled hazelnuts (Jacobs, 1992), and on American crows (*Corvus brachyrhynchos*), which are more likely to store black walnuts, which take longer to handle compared to English walnuts (Cristol, 2001). Although handling time is rarely considered in discounting tasks, work has demonstrated that it does influence how tamarins (*Saguinus oedipus*) and marmosets (*Callithrix jacchus*) devalue rewards, both being less likely to choose the LL reward when handling time is increased (Rosati et al., 2006).

When we increased the handling time for both food types by the same amount, the foragers in our model became less likely to eat SS items. This is because the SS food contains less energy, so the animal faces a greater energy loss: gain ratio when eating this food (Krebs et al., 1977). The predicted reduction in the probability of eating SS items is most prominent in the laboratory scenarios, when the forager faces a low predation risk while searching and a high probability of finding another food item. Given that foragers can expect to encounter LL food items frequently under these conditions, they can increase their immediate energy reserves more quickly if they avoid spending time handling the less profitable SS items.

Foragers never rejected encountered food items in our simulated urban and rural environments, where food is scarcer. Foragers did not start to reject the LL item with an increase in handling time, even when it was more abundant in the environment. We would anticipate that foragers could afford to be more selective in these circumstances, because after rejecting a food item, they are likely to quickly find another item. It is therefore likely that other remaining risks (*e.g.*, predation) prevent the animal from rejecting the LL item in these conditions.

Our prediction that foragers should only reject food in laboratory scenarios is further supported by classic work on great tits (*Parus major*), which reject less profitable prey items when they encounter more profitable prey items at high rates (Krebs et al., 1977). Similarly, three-spined sticklebacks (*Gasterosteus aculeatus*) are most likely to attack the first prey they encounter when food is less abundant (Ioannou et al., 2008). This empirical work aligns with models of partial preferences, where animals may reject food items, but not in an all-or-none approach (McNamara & Houston, 1987) as suggested by the Optimal Diet Model.

In the model of McNamara & Houston (1987), animals following a strategy that minimises the probability of starvation are the least likely to reject a food item when their energy reserves are low, there is a short time left until the end of the foraging period (at which point the animal must reach a critical amount of energy), and their probability of finding another food item is low. Like the foragers in our model, empirical work (Krebs et al., 1977; Ioannou et al., 2008) demonstrates that animals only rejected encountered food items when the probability of finding another food item was high. Although the LL starts as the more profitable food type, when handling time for only this food type is increased (while holding *h*_SS_ constant), the SS food item becomes the more profitable food item, explaining why the LL item is rejected at higher handling times in our simulated laboratory scenarios.

Partial preferences were predicted by our model, possibly because foragers must utilise encountered food for energy reserves stored both in their bodies and in caches for winter. Food-hoarding animals may show different preferences based on whether they are making decisions with more immediate *vs.* future outcomes (*e.g.*, Steele et al., 2001b). In our model, foragers are never likely to exclude a food type completely from their diet because, for example, even when the handling time to the LL item is long and foragers may prefer to eat SS, the LL food retains utility for caching.

### Predation Risk

Under the optimal strategy in simulated natural conditions, foragers never reject encountered food, likely because searching entails a high predation risk. A forager will only rest when they have an adequate amount of energy reserves and thereby face a low risk of starvation. This is in line with other models that show that as an animal’s energy reserves increase, it should take fewer risks that could lead to predation (*e.g.*, McNamara, 1990), and empirical work showing that free-living squirrels always eat or cache the items they encounter (Preston & Jacobs, 2009). In particular, our model predicts that a food-storing animal should only reject food in favour of resting in extreme circumstances, such as nearing or reaching a limit on the amount of energy cached – a state that is unlikely to be observed in wild animals.

Foragers only reject food items under laboratory conditions in our model, where the risk of predation while searching is low. This prediction aligns with previous work on beetles (*Harpalus affinis*), salmon (*Salmo salar*), and bees (*Bombus terrestris*), which all become less selective towards food items when exposed to predator cues (Charalabidis et al., 2017; Metcalfe et al., 1987; Wang et al., 2013b; although see Leaver & Daly, 2003). Although this reduction in choosiness can cause animals to eat lower-value food items they may normally reject, this trade-off can reduce predation risk, where time spent assessing food items can instead be invested into vigilance (Charalabidis et al., 2017). Similarly, in our model, foragers avoid rejecting less profitable items (*e.g.*, LL items as handling time for this item increases) in urban and rural environments, where searching for an alternative item would entail a high chance of death.

Notably, when the handling time for the LL food is increased while that for the SS is kept constant, foragers in our model start to reject LL items in laboratory scenarios. In contrast, when the handling time for both food types is increased, foragers in our model start to reject SS items. These predictions are reminiscent of how animals behave in laboratory discounting tasks, where individuals choose the LL reward when it is freely available, but opt for the SS reward as the delay to LL increases and show a preference reversal when the delays to both rewards are increased (Ainslie, 1974). We can question whether laboratory foraging scenarios with negligible predation risk while searching and guaranteed food are realistic. Like Krebs et al.’s (1977) great tits, which simply had to wait for the conveyor belt to bring them a food item, with no risk of predation, animals in traditional discounting tasks always receive either an SS or an LL item, and there is no risk while enduring the wait for this reward.

There has been very little research on how predation risk influences discounting. Rats (*Rattus norvegicus*) tolerate longer delays for LL rewards when the food item is in an enclosed *vs.* an open maze, reducing perceived predation risk (Feeney & Roberts, 2008), suggesting that animals should prioritise more immediate outcomes in these circumstances. Similarly, Canada jays (*Perisoreus canadensis*) are less likely to opt for a larger reward when they must travel further into a tube to collect it, making it harder for them to escape (Waite, 2001); although, it is important to note that delay and effort may have been confounded with predation risk in this study. In our model, when faced with a high risk of predation, foragers are predicted to increase their chances of survival by accepting all encountered food items, avoiding searching for a new item and thereby reducing their exposure to risk.

### Collection Risk

Our model predicts that foragers should reject the LL food with long handling times at lower rates in the laboratory scenario with pilfering, compared to the laboratory scenario without pilfering. A forager that is consistently losing stored items should cache encountered items more frequently (Huang et al., 2011; Lucas & Zleliniski, 1998; but see Lucas & Walter, 1991) to compensate for their losses, ensuring they survive winter and leading to lower rejection rates. Our model also predicts that foragers are most likely to cache the LL food when pilferage risk is high, suggesting that, despite their high handling times, these items are valuable for counteracting frequent cache loss to reduce the risk of starvation when winter arrives.

Conversely, in laboratory conditions without pilfering, stored items remain more secure. Therefore, animals can be more selective, rejecting LL items with long handling times and searching for lower-cost items, without jeopardising their survival. However, in rural environments where pilferage risk is similarly low, foragers never reject any food items. These contrasting results suggest that other risks need to be suitably high (*e.g.*, predation risk) for foragers to show reduced selectivity in our model.

Foragers may also fail to show increased selectivity in rural environments with reduced pilferage due to food availability. The steepest increase in rejection with handling time is seen in our model when LL foods are more abundant in the environment. Foragers are predicted to reject the SS food at relatively high rates in the laboratory condition with pilfering, when the handling time for LL items is low, and LL items are highly abundant in the environment. When the handling time for LL items is very low, the forager receives a larger benefit from eating or caching these items than they do from SS items, and the high chance of finding an LL item means the forager can afford to reject SS items without risk of starvation. As the handling time for LL items increases, the forager almost exclusively relies on caching, rather than eating these items, meaning they can no longer reject SS items and instead must consume them to increase their immediate energy reserves.

When collection risk is factored into discounting studies, we typically see steeper devaluation of delayed rewards. This is in line with the Collection Risk Hypothesis, which predicts that animals should prioritise more immediate rewards when interruptions may prevent them from obtaining a delayed reward (Stevens & Stephens, 2010). For example, Eurasian jays (*Garrulus glandarius*) are more likely to choose an SS reward in the presence of conspecifics (Miller et al., 2023), bonobos (*Pan paniscus*) generally show a greater preference for SS outcomes when rewards are uncertain due to the presence of an unreliable experimenter (Stevens et al., 2011), and chicks (*Gallus gallus domesticus*) trained under direct and perceived competition are more likely to opt for the SS reward (Amita et al., 2010). Similarly, foragers in our model always accept encountered items in urban and rural environments, even if the items have low profitability, to reduce cache loss; if the forager searches for another item in these environments, it is likely to take them an extended amount of time to find an alternative due to lower food availability, potentially resulting in a higher chance of pilferage, a form of collection risk, where the risk of stored food being taken by a competitor scales with time.

### The Effect of Hoarding

When caching is switched off in our model, we see similar patterns of rejection dependent on the simulated environment. Foragers seldom reject food items in natural environments but show steep rejection for the LL item with an increase in handling time to the LL item only, and for the SS item with an increase in handling time to both food items in laboratory environments. Unsurprisingly, this pattern is the same between urban and rural environments and laboratory environments with and without pilfering respectively, highlighting that pilfering does not influence these results due to a lack of value in hoarding food. As expected, non-hoarding foragers show very high rates of rejection for both food items with these increases in handling time. For a non-hoarding forager, food items no longer retain utility for caching despite their high handling times, likely producing this result.

Overall, predictions from our model suggest that high levels of rejection for the LL reward with an associated delay in discounting tasks may be an artefact of artificial laboratory conditions, where food is guaranteed, and there is little collection or predation risk. Under these conditions in our model, the forager can be more selective, because there is less risk of dying from starvation, from not meeting the target amount of energy cached when winter starts, or from predation. As the handling time for LL food or for both foods is increased, the forager may sometimes reject LL items or SS items, respectively, while maintaining a high chance of survival. Our predictions suggest that high rejection of LL food in laboratory conditions with an increasing delay is not maladaptive, but it does not align with natural foraging behaviour, because of a mismatch between the testing environment and the environment in which animals have evolved to make decisions.

